# Comparison of the therapeutic effects of hUC-MSC intravenous delivery and intraperitoneal administration of MSCs encapsulated in alginate capsules for the treatment of rat liver cirrhosis

**DOI:** 10.1101/2021.04.26.441497

**Authors:** S. Rymar, P. Pikus, P. Buchek, N. Shuvalova, Ia. Pokholenko, D. Irodov, V. Kordium

**Author notes:** **Competing interests:** The authors have declared that no competing interests exist.

## Abstract

Mesenchymal stem cells are the most promising regenerative medicine tool for the treatment of various diseases, including liver disease, although the exact mechanism of their therapeutic action remains unclear. It was found that MSCs are captured by the lungs after systemic transplantation, quickly disappear, and are not detected at the site of injury, but at the same time exhibit an obvious therapeutic effect. Comparison of the MSC efficiency depending on the route of their administration may shed light on the mechanisms involved in the implementation of MSC therapeutic potential. In this work, we compared the therapeutic effects of human umbilical cord MSCs (hUC-MSCs) administered systemically and intraperitoneally in the form of MSCs encapsulated in alginate capsules in a CCl_4_-induced model of liver cirrhosis in rats. Our study showed that both treatments resulted in liver recovery. MSC transplantation by two different routes led to a decrease in collagen deposition, the disappearance of the fibrous area by the 13^th^ week, and normalization of the morphometric parameters of liver parenchyma cells. The expression of some genes (EGF, alpha SMA, GFAP) which is activated in liver injury, decreased to the level observed in negative control animals. However, a detailed study of liver recovery in dynamics showed that encapsulated MSCs led to faster normalization in several parameters of the liver tissue. Our results showed that human umbilical cord MSCs effectively exhibit their therapeutic properties when using both methods of transplantation, however, intraperitoneal administration of encapsulated MSCs accelerated the process of liver regeneration.

## Introduction

Liver cirrhosis is a serious illness that most often results in death. Currently, the liver transplantation was the only method of treating cirrhosis. However, as an alternative, the possibilities of cell therapy are being studied, and in particular the use of mesenchymal stem cells (MSCs) of various origins.

Studies of the effect of MSCs from various tissue sources on liver diseases carried out in animals with experimental liver damage showed a therapeutic effect, and clinical trials on patients with cirrhosis demonstrated the safety of transplantation of MSCs and liver improvement without long-term effect. How MSCs affect the processes occurring in the injured liver have been considered in numerous reviews [1-11]. However, despite this large number of studies on liver recovery through transplantation of MSCs our understanding of the role of MSCs in liver failure is still incomplete. One of the main mechanisms responsible for the therapeutic properties of MSCs is their immunomodulating properties, which are mainly caused by the secretion of a large number of cytokines, chemokines, and growth factors, due to which they interact with cells of the immune system [12-20].

Numerous studies demonstrate different features of the MSC interaction with immune cells, in particular, the possibility of direct cell contacts, which were studied in detail *in vitro* [21], modulation of MSC exosomes, which are the mediator of cell-to-cell communication [22-24]. Another way to achieve immunomodulation is the activation of macrophages [25] and monocytes as a result of phagocytosis of MSCs [26, 27].

Thus, today the question of the mechanisms providing the therapeutic effects of MSCs remains open. The use of different routes of MSCs transplantation makes it possible to compare the therapeutic consequences of such an introduction of MSCs and to relate them to the implementation of different immunomodulation mechanisms. This study aimed to compare the therapeutic effect of intravenous administration of human umbilical cord MSCs with the effect of intraperitoneal administration of MSCs encapsulated in alginate capsules for the treatment of rat liver cirrhosis induced by CCl_4_.

## Materials and methods

### Induction of liver cirrhosis in rats

Experiments were performed on male Wistar rats, 2-3 months of age. All animal experiments were carried out according to the Resolution of the Presidium of the Methodological recommendations bioethics examination of scientific studies conducted on animals (National Bioethics Committee at the Presidium of NAS of Ukraine, 2006). The studies were approved by the Institute of Molecular Biology and Genetics of the NAS of Ukraine, that they do not conflict with the bioethical issues. The rats were kept in groups in standard plastic cages containing sawdust. During the experiment, the rats were divided into two groups: the negative control (n=15) and the experimental group (n=30). Liver injury was caused by the intraperitoneal injections of a CCl_4_ solution in olive oil (1:1) twice weekly as described in Table 1. Rats were weighed before each injection to calculate the required dose of a CCl_4_. The control animals were injected intravenously with a mixture of saline and olive oil (1:1).

**Table 1.** Scheme of CCl_4_ induction of rats liver injury.

| Animal groups | 2 weeks CCl<sub>4</sub> administration | 3-4 weeks CCl<sub>4</sub> administration | 5-13 weeks CCl<sub>4</sub> administration |
| --- | --- | --- | --- |
| **Negative control** (norm) (n=15) | 0,1 ml mixture (olive oil: saline)/100g rat weight | 0,075 ml mixture (olive oil: saline)/100g rat weight | 0,05 ml mixture (olive oil: saline)/100g rat weight |
| **Experimental group** (cirrhosis) (n=30) | 0,1 ml mixture (CCl<sub>4</sub>: olive oil)/100g rat weight | 0,075 ml mixture (CCl<sub>4</sub>: olive oil)/100g rat weight | 0,05 ml mixture (CCl<sub>4</sub>: olive oil)/100g rat weight |

### Histology

For histological analysis and RNA isolation to study the dynamics of liver regeneration, liver samples were recovered by partial hepatectomy (3-4% of the total liver volume) under anesthesia. Liver samples were subsequently fixed in 10% neutral buffered formalin, embedded with paraffin, sectioned at a thickness of 5-6 μm, and then stained with hematoxylin and eosin (H&E) to study the morphometric parameters of liver tissue [28]. The area of fibrosis was expressed as a percentage of the total liver area [29]. To determine this area, liver sections were stained according to Van Gieson and analyzed 15 fields of liver sections per animal. Each field was obtained at a magnification of 10×10 (microscope Leica and digital camera Sigeta). Collagen area and total field area were determined using the ImageJ software (version 1.52a). The area subtraction of the vascular lumen from the total area of the field gave the final calculation of the net area of fibrosis.

### Isolation and characterization of hUC-MSCs

Human umbilical cords from both sexes were collected from full-term births after normal vaginal delivery with informed consent using the guidelines approved by the Institutional Ethics Committee (IEC) at the Municipal Maternity Hospital №5, Kyiv, Ukraine. MSCs were isolated from a human umbilical cord after removal of the umbilical blood vessel following the explant method [30]. MSCs were cultivated in α-MEM medium (HyClone, Thermo Scientific) containing 10% fetal bovine serum (HyClone, Thermo Scientific), 200 U/ml penicillin, and 200 µg/ml streptomycin. After the second passage, MSCs were evaluated for surface marker expression and differentiation capacity. The adipogenic, osteogenic, and chondrogenic differentiation potentials of MSCs were tested using the StemPro® Differentiation Kits (Gibco) (Fig.1A - C). We used Alcian blue staining (1% Alcian blue (Sigma-Aldrich) in 3% glacial acetic acid solution) to observe chondrogenic differentiation. For osteogenic differentiation, the cells were analyzed by staining with Alizarin Red S (Sigma-Aldrich) and for adipogenic differentiation, the cells were stained with Oil Red O (Sigma-Aldrich). Surface markers of MSCs were assessed by flow cytometry with FACS Aria (Becton Dickinson Lakes, NJ) using CD90 FITC-A, CD73 APC-A, CD105 PerCP – Cy5-5-A, CD45 FITC – A, CD34 APC – A. (Fig. 1 D)

**Fig 1.**
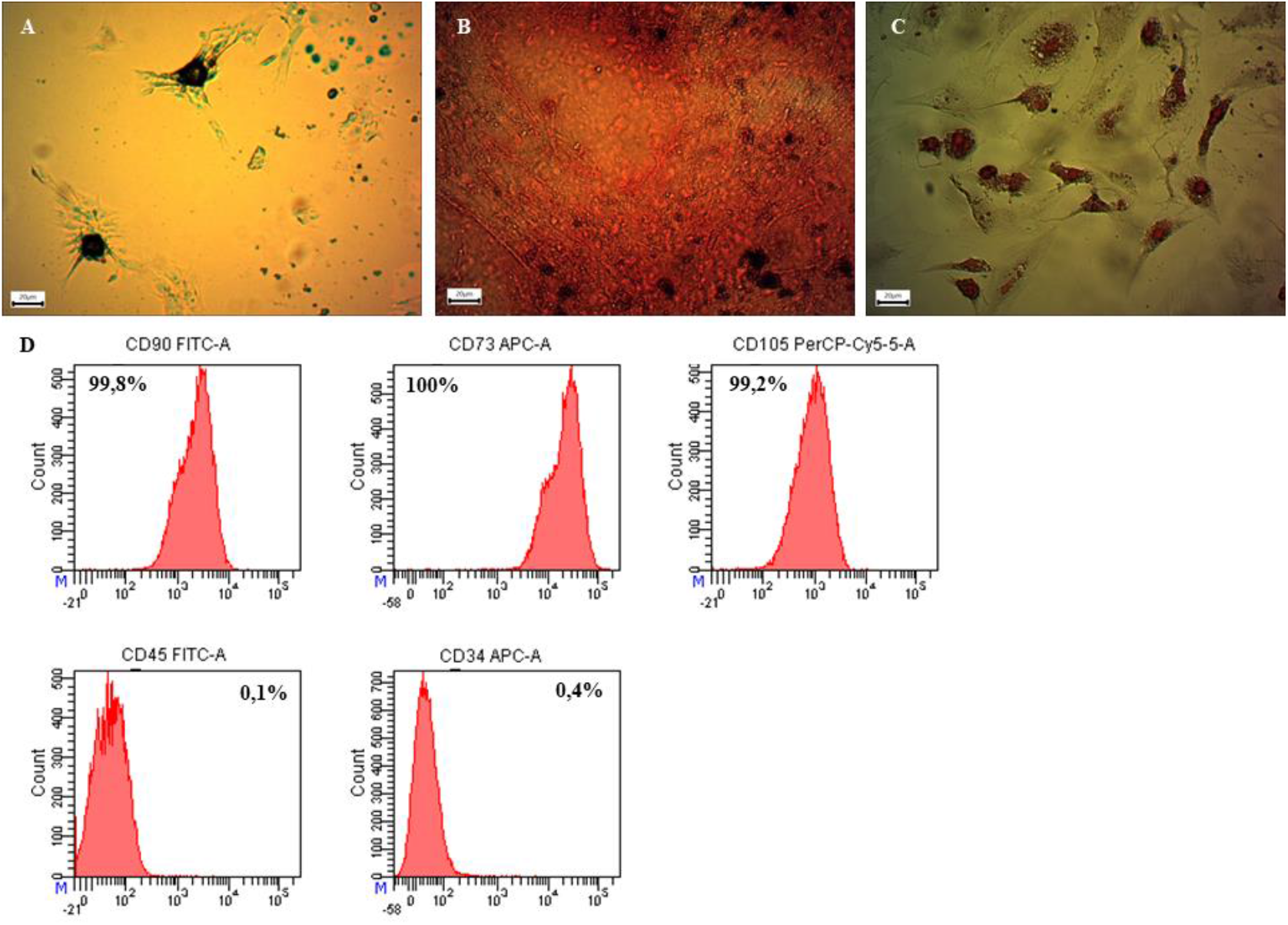
The characteristics of hUC-MSCs: **A** - chondrogenic differentiation was detected by Alcian blue staining; **B** - osteogenic differentiation was detected by Alizarin red S staining; **C** - adipogenic differentiation was detected by Oil Red O staining; **D** – expression of MSC surface markers from human umbilical cord. Scale bar 20 μm.

### Analysis of gene expression in rats

Total RNA was isolated from liver samples using the NucleoSpin RNA Kit (Macherey-Nagel) according to the manufacturer’s instructions. For cDNA synthesis, 1 μg RNA was used as a template for reverse transcription (RT) using the RevertAid H Minus First Strand cDNA Kit (Thermo Scientific). Sequence of primers used in this study is shown in Table 2. The mRNA levels of the target genes were normalized against the mRNA level of GAPDH. ImageLab program was used for densitometry analysis of electrophoresis data.

**Table 2.**
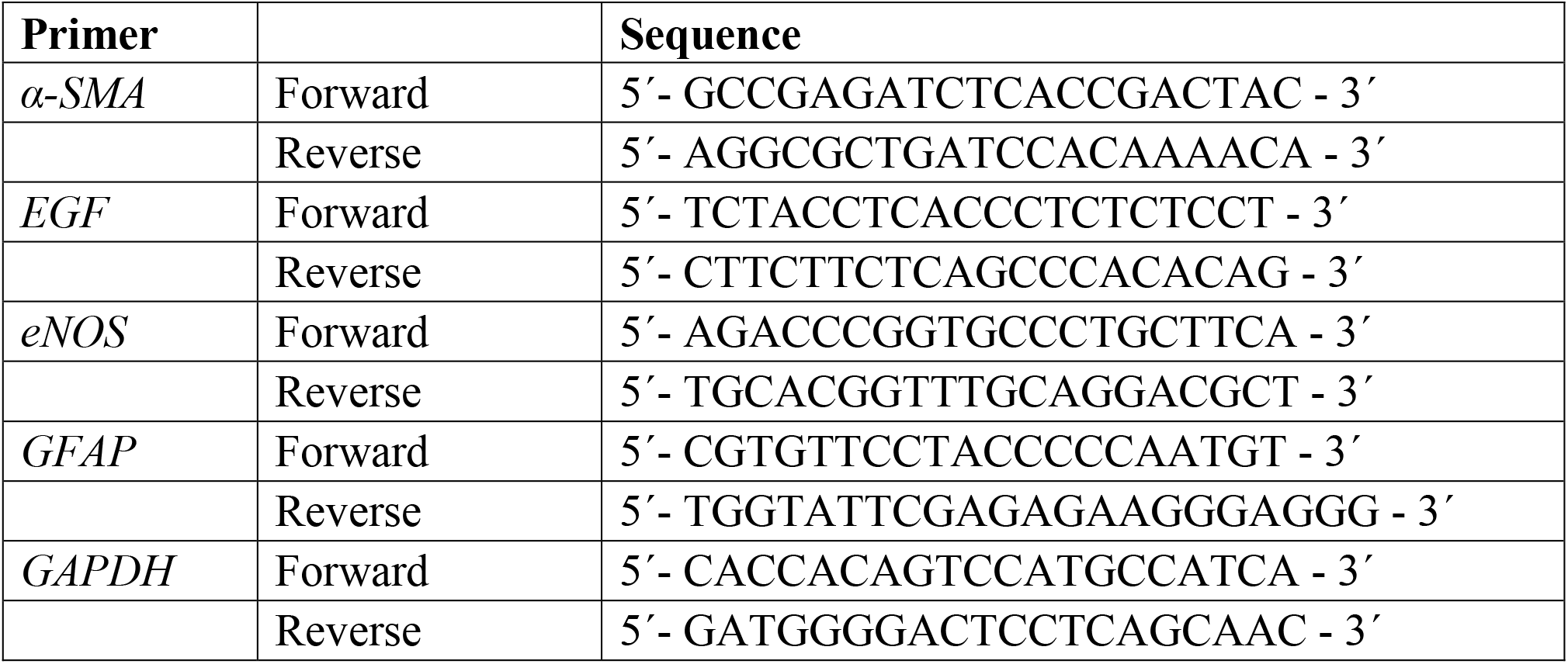
List of primers used.

### Encapsulation of hUC-MSCs

The MSCs of the second passage were trypsinized, collected by centrifugation, washed three times with PBS. MSCs were resuspended in 150 μl saline and encapsulation occurred by mixing the MSCs with 1.7% sodium alginate (Sigma) in saline (10^7^ cells/ml). Using a 29g syringe, the cell suspension was added dropwise to a 102 mM CaCl_2_ solution [31]. The capsules were incubated for 30 min, washed three times with PBS, immersed in a 0.1% solution of poly-L-ornithine, stirred for 30 min, and washed with PBS three times more (Fig. 2).

**Fig 2.**
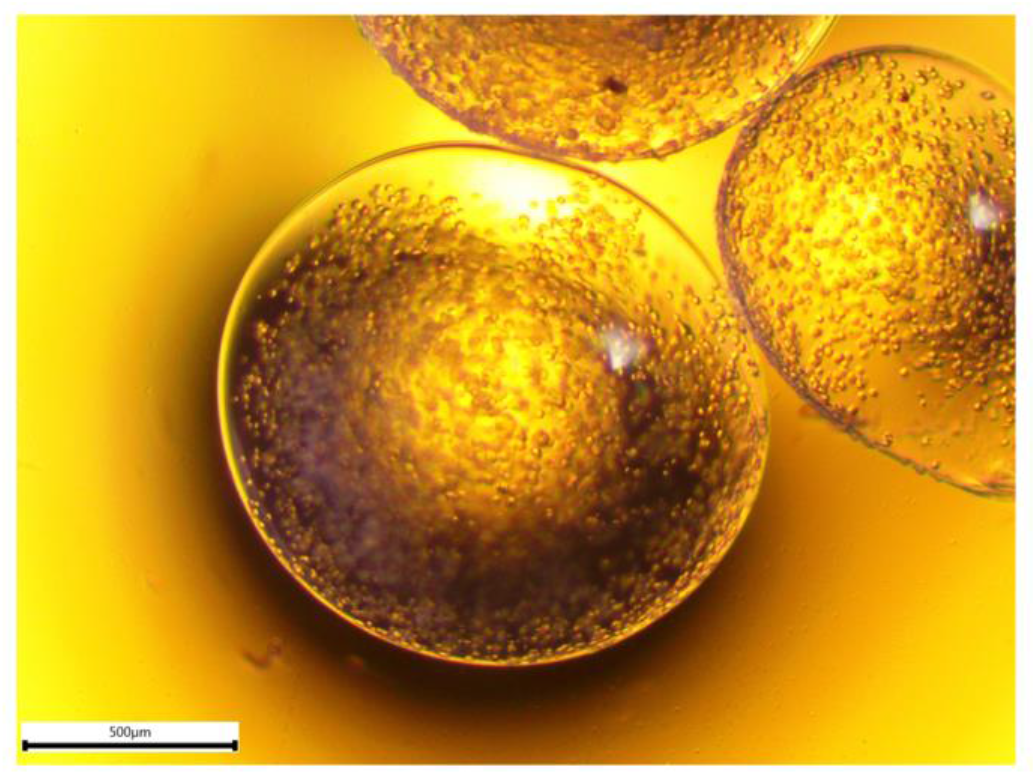
Encapsulated MSCs in alginate capsules. Scale bar 500μm.

### Transplantation of hUC-MSCs

After 13 weeks of CCl_4_ administration to the rats was stopped (0 point). The animals were divided into 4 groups of 6 animals each. The animals of the first group received 150mkl suspension of hUC-MSCs (6 × 10^6^ cells/kg) in the tail vein, while the animals of the second group were intraperitoneally injected with MSCs encapsulated in the alginate capsules in the same amount as in the first group. The animals of the third group received 150mkl PBS in the tail vein (sham control-PBS), the fourth group was injected intraperitoneally with “empty” alginate capsules (sham control-”empty” capsules).

### Statistical analysis

Statistical significances of PCR and morphometric parameters of animal liver were determined with the use GraphPad Prism software, Version 8.0.1. Comparisons of negative control and experimental group (cirrhosis) were performed using unpaired two-tailed Student’s t-tests. Comparisons involving multiple groups were evaluated via one-way analysis of variance (ANOVA) followed by Tukey’s test. For all tests, p < 0.05 was considered significant.

## Results

### The development of liver injury induced by the injections of CCl_4_

Intraperitoneal injections of CCl_4_ led to signs of rat liver injury. Histological study of the liver for 13 weeks with systemic administration of CCl_4_ showed increasing changes in the liver parenchyma of experimental animals as compared to the negative control. The area occupied by fibrous tissue gradually increased (Fig.3B, E, Table 3). H&E staining of liver sections showed that with an increase in the number of intraperitoneal injections of CCl_4_ the morphology of hepatocytes changed; the nuclear-cytoplasmic ratio decreased; the number of binuclear cells decreased; the number of nuclear-free cells increased 11-fold (Table 3). By the 13^th^ week, the area of fibrous tissue was more than 6 times larger compared to the negative control (Fig.3F, Table 3) and collagen accumulation led to the appearance of partitions between themselves dividing the parenchyma into separate fragments. This indicates the formation of pseudoglobules, which are a sign of liver cirrhosis (Fig.3C).

A study of the EGF gene expression (epidermal growth factor), which, as is known, almost unexpressed in a healthy liver and begins to be expressed with the development of fibrosis and is intensively expressed in a cirrhotic liver [32] showed a sharp increase in the level of RNA expression (Fig. 4A).

**Table 3.** Morphometric parameters of rat liver after CCl_4_ induction. Negative control – intravenous administration of olive oil and saline; liver fibrosis – 8 weeks after intraperitoneal administration of CCl_4_ to rats; liver cirrhosis – 13 weeks after intraperitoneal administration of CCl_4_ to rats. Results are expressed as mean±SEM. P values were calculated by ANOVA with Tukey’s test.

| Morphometric parameters | Negative control (n=15) | Experimental group (n=30) |  |  |  |
| --- | --- | --- | --- | --- | --- |
|  |  | Fibrosis, 8 weeks |  | Cirrhosis, 13 weeks |  |
|  |  | Mean | P-value<sup>*</sup> | Mean | P-value<sup>#</sup> |
| **Collagen accumulation** | 0,06±0,007 | 0,24±0,02 | **0,0001** | 0,39±0,02 | **<0,0001** |
| **Hepatocyte area, mkm<sup>2</sup>** | 155±4,65 | 123,2±6,01 | **0,0008** | 108±6,5 | **<0,0001** |
| **Hepatocyte nucleus area, mkm<sup>2</sup>** | 44,7±5,5 | 31,3±7,7 | **<0,0001** | 23,1±6,5 | **<0,0001** |
| **Nucleocytoplasmic ratio** | 0,43±0,009 | 0,35±0,009 | **<0,0001** | 0,25±0,009 | **<0,0001** |
| **Binuclear hepatocytes** | 8,2±0,64 | - | - | 2±0,72 | **<0,0001** |
| **Nuclear-free hepatocytes** | 0,57±0,29 | - | - | 5,4±0,64 | **<0,0001** |

<sup>\*</sup> p<0.05 – negative control vs. fibrosis
<sup>#</sup> p<0.05 – negative control vs. liver cirrhosis

**Fig 3.**
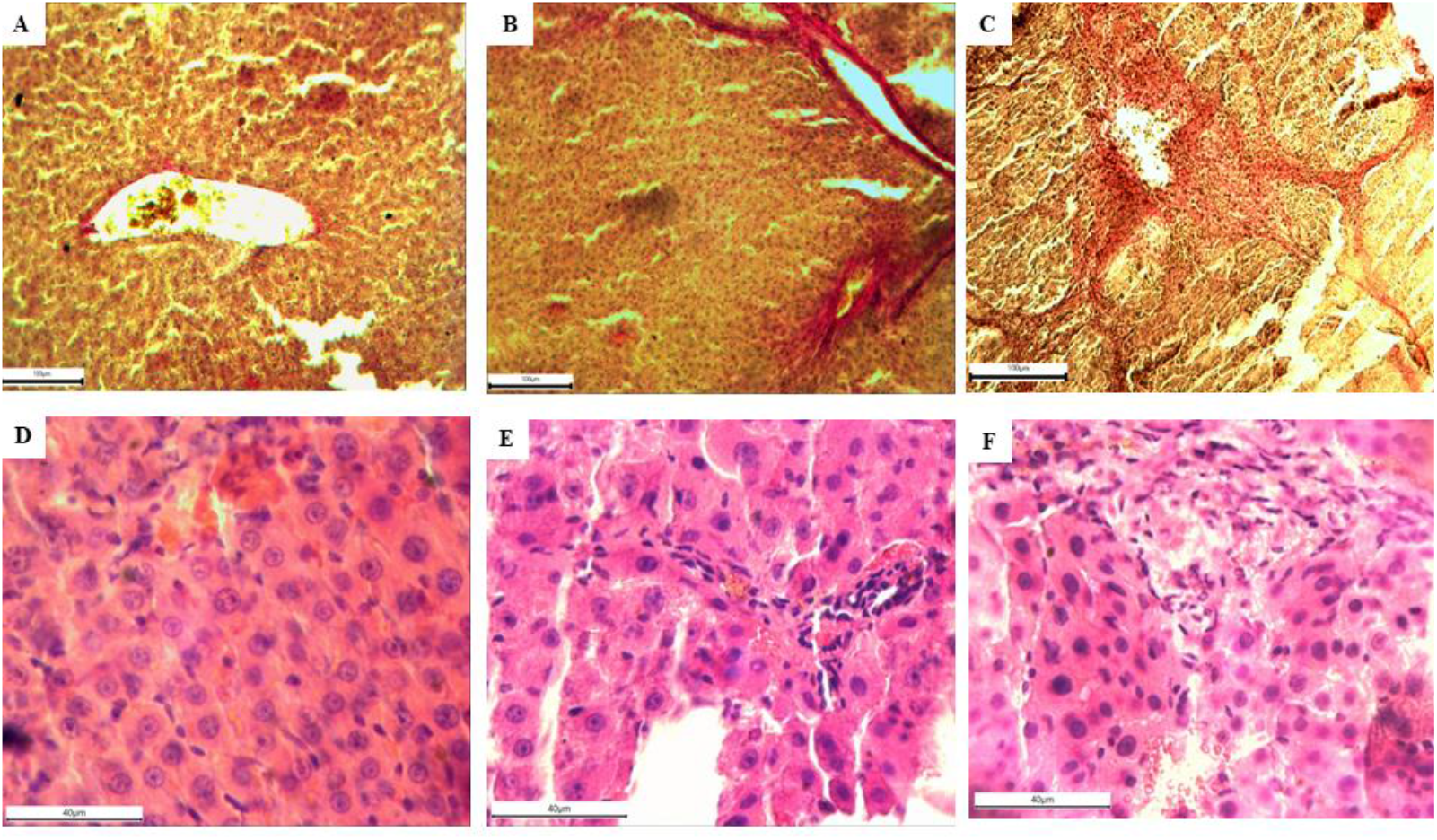
Histological analysis of liver tissue sections after CCl _4_ induction of liver fibrosis/cirrhosis in rats. Van Gieson staining and H&E staining of rat liver sections: **A, D** – negative control (intravenous injection of mixture (saline : olive oil)); **B, E** - liver sections from rats administered intraperitoneally with CCl_4_ for 8 weeks; **C, F** - liver sections from rats administered intraperitoneally with CCl_4_ for 13 weeks. Scale bar: **A - C** - 100 μm, **D – F** - 40 μm

**Fig 4.**
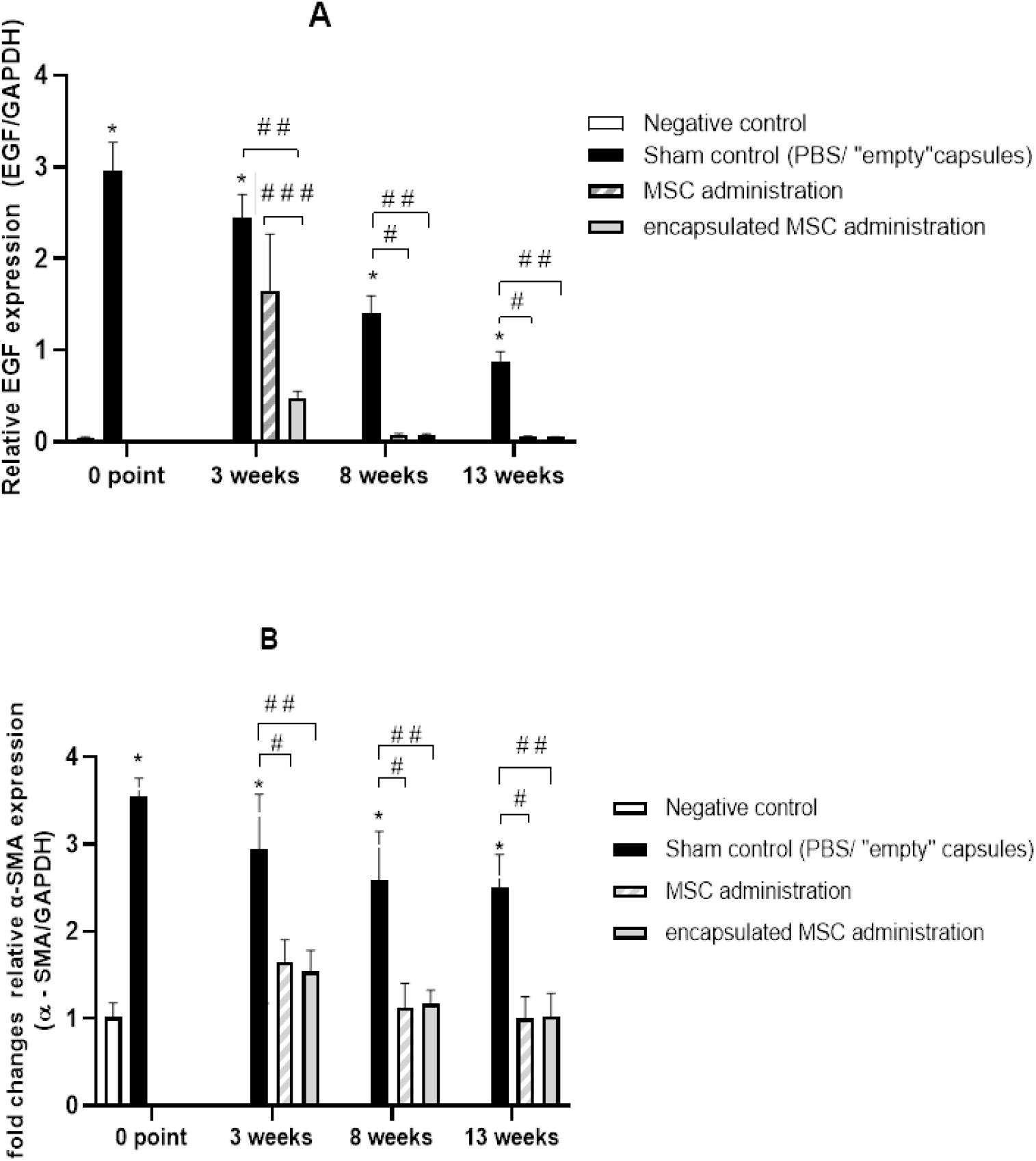
The expression of EGF (A) and α-SMA (B) genes: 0 point - 13 weeks after intraperitoneal administration of CCl_4_ to rats; 3, 8 and 13 weeks - after intravenous administration of MSCs and intraperitoneal administration of encapsulated MSCs to rats with liver cirrhosis. Results are expressed as mean±SEM. P values were calculated by ANOVA with Tukey’s test. There were no significant differences between the (sham control-PBS) and (sham control-«empty» capsules) – 0,93 < p < 0,99. ^*^ p<0.01 – negative control vs. sham control ^#^ p<0.01 – sham control vs. MSC administration ^##^ p<0.01 – sham control vs. encapsulated MSC administration ^###^ p<0.01 – MSC administration vs. encapsulated MSC administration

The α-SMA gene (alpha-smooth muscle actin) is a reliable marker of hepatic stellate cell activation during the development of fibrosis and accompanies an increase in the number of myofibroblasts [33, 34] and the expression level of this gene increased too (Fig. 4B).

GFAP (glial fibrillary acidic protein) expression, which is known to be associated with early activation markers of stellate cells [35, 36], increased 2.5-fold (Fig. 5A). Expression of the eNOS gene (endothelial nitric oxide synthase) decreased slightly (p<0.01) (Fig. 5B), in contrast to significant decrease of the eNOS expression that was shown in the CCl_4_ -induced model of liver injury in mice [37].

**Fig 5.**
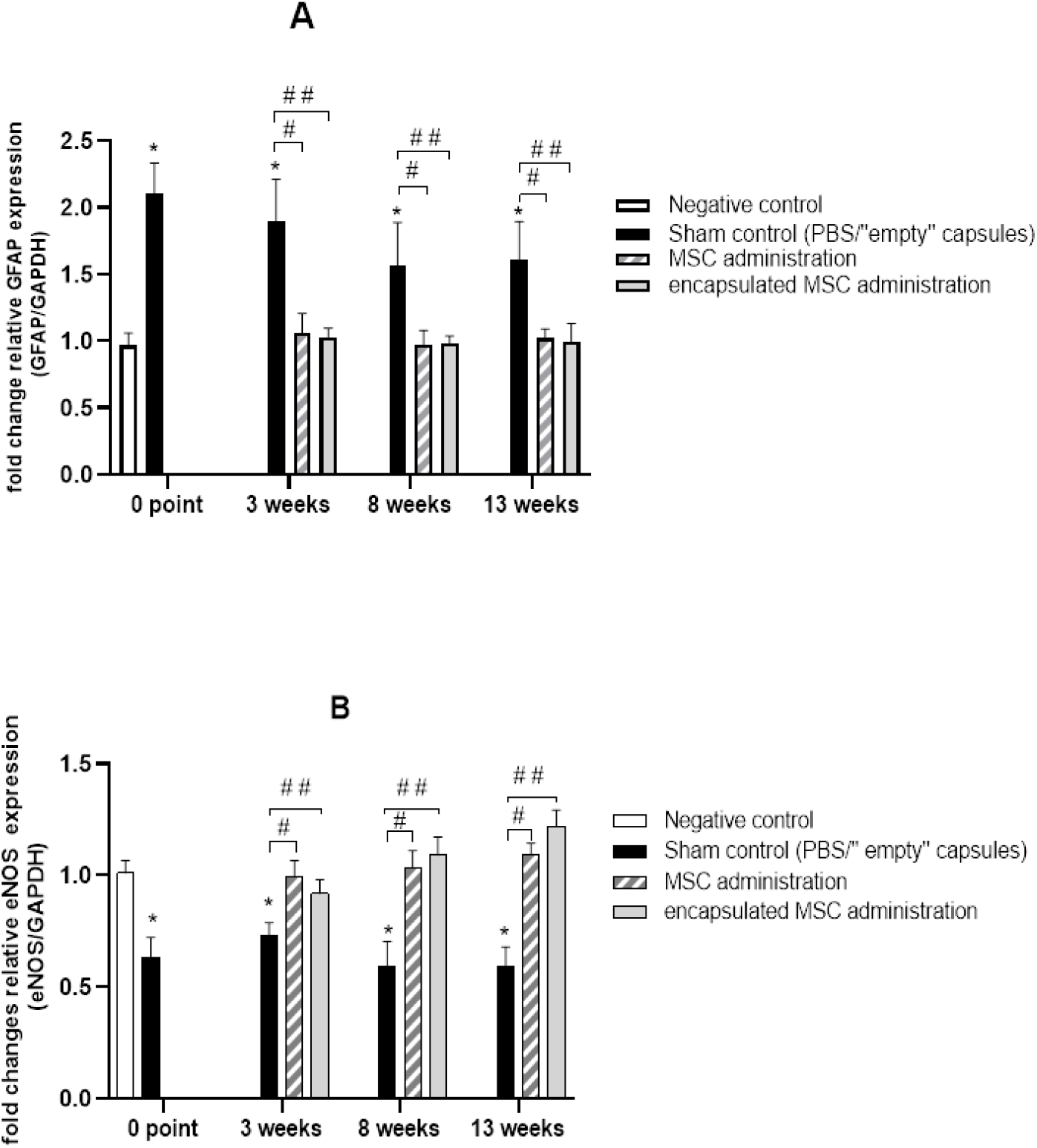
The expression of GFAP (A) and eNOS (B) genes: 0 point - 13 weeks after intraperitoneal administration of CCl_4_ to rats; 3, 8 and 13 weeks - after intravenous administration of MSCs and intraperitoneal administration of encapsulated MSCs to rats with liver cirrhosis. Results are expressed as mean±SEM. P values were calculated by ANOVA with Tukey’s test. There were no significant differences between the (sham control-PBS) and (sham control-”empty” capsules) – 0,75 < p < 0,99. ^*^ p<0.01 – negative control vs. sham control ^#^ p<0.01 – sham control vs. MSC administration ^##^ p<0.01 – sham control vs. encapsulated MSC administration

### Effect of MSC transplantation on the rat liver cirrhosis

After the thirteen weeks of CCl_4_ injection, when signs of cirrhosis appeared, CCl_4_ administration was discontinued and analysis of changes in the structure of the liver parenchyma and the expression of the selected genes was determined after 3, 8, and 13 weeks after transplantation of human umbilical cord MSCs. Dynamics of changes in the fibrosis level occurring in rat liver after transplantation of MSCs (Fig.6) demonstrates that the injured liver underwent a regeneration process.

The fibrosis area after MSC transplantation gradually decreased within three weeks in particularly, by 24% after systemic administration of MSCs and by 43% after the introduction of encapsulated MSCs (Fig. 7B). Morphometric parameters of the liver tissue (Table 4) also indicate repair processes in the liver in animals that received MSCs both in the form of a cell suspension and as cells encapsulated in alginate capsules. Parameters of hepatocytes returned to the values observed in control animals, the number of dividing cells increased, and the number of nuclear-free cells decreased. By the thirteenth week, the morphometric indicators of the experimental animal liver were significantly not different from those that were not injected with CCl_4_. Liver recovery in animals that were exposed to CCl_4_ but did not receive MSCs was extremely slow and almost not observed even after 13 weeks (Fig. 7A). Although liver recovery in rats with induced cirrhosis, when the toxic factor disappears, occurs from 12 to 24 weeks if the cirrhosis is micronodular [24].

**Table 4.**
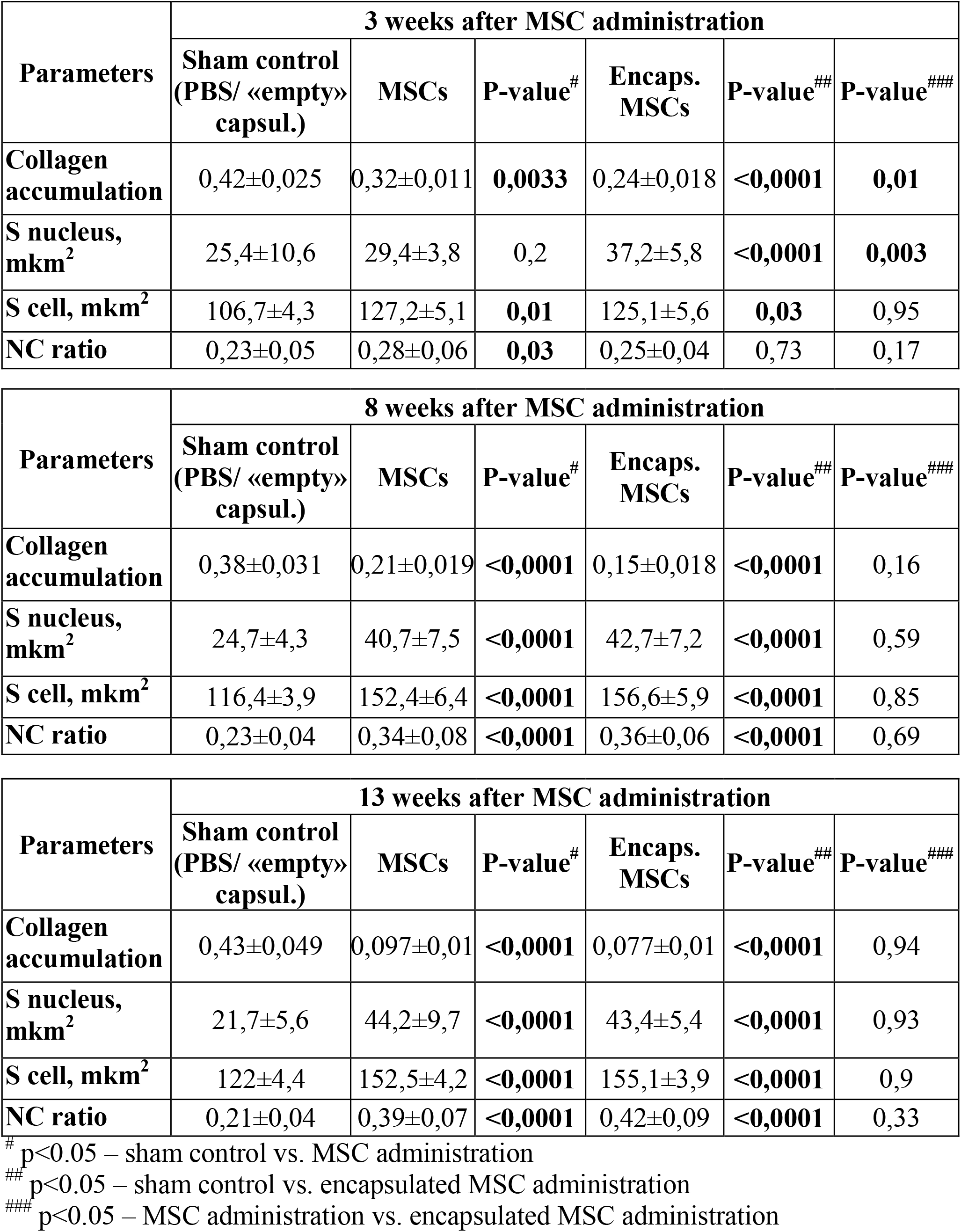
Morphometric analysis of rat liver tissue 3, 8, and 13 weeks after intravenous administration of MSCs and intraperitoneal administration of encapsulated MSCs to rats with liver cirrhosis. Results are expressed as mean±SEM. P values were calculated by ANOVA with Tukey’s test. There were no significant differences between the sham control-PBS and sham control-«empty» capsules - 0,73<p<0,99

| Parameters | 3 weeks after MSC administration |  |  |  |  |  |
| --- | --- | --- | --- | --- | --- | --- |
|  | Sham control<br>(PBS/ «empty»<br>capsul.) | MSCs | P-value<sup>#</sup> | Encaps.<br>MSCs | P-value<sup>##</sup> | P-value<sup>###</sup> |
| **Collagen**<br>**accumulation** | 0,42±0,025 | 0,32±0,011 | **0,0033** | 0,24±0,018 | **<0,0001** | **0,01** |
| **S nucleus,**<br>**mkm<sup>2</sup>** | 25,4±10,6 | 29,4±3,8 | 0,2 | 37,2±5,8 | **<0,0001** | **0,003** |
| **S cell, mkm<sup>2</sup>** | 106,7±4,3 | 127,2±5,1 | **0,01** | 125,1±5,6 | **0,03** | 0,95 |
| **NC ratio** | 0,23±0,05 | 0,28±0,06 | **0,03** | 0,25±0,04 | 0,73 | 0,17 |

| Parameters | 8 weeks after MSC administration |  |  |  |  |  |
| --- | --- | --- | --- | --- | --- | --- |
|  | Sham control<br>(PBS/ «empty»<br>capsul.) | MSCs | P-value<sup>#</sup> | Encaps.<br>MSCs | P-value<sup>##</sup> | P-value<sup>###</sup> |
| **Collagen**<br>**accumulation** | 0,38±0,031 | 0,21±0,019 | **<0,0001** | 0,15±0,018 | **<0,0001** | 0,16 |
| **S nucleus,**<br>**mkm<sup>2</sup>** | 24,7±4,3 | 40,7±7,5 | **<0,0001** | 42,7±7,2 | **<0,0001** | 0,59 |
| **S cell, mkm<sup>2</sup>** | 116,4±3,9 | 152,4±6,4 | **<0,0001** | 156,6±5,9 | **<0,0001** | 0,85 |
| **NC ratio** | 0,23±0,04 | 0,34±0,08 | **<0,0001** | 0,36±0,06 | **<0,0001** | 0,69 |

| Parameters | 13 weeks after MSC administration |  |  |  |  |  |
| --- | --- | --- | --- | --- | --- | --- |
|  | Sham control<br>(PBS/ «empty»<br>capsul.) | MSCs | P-value<sup>#</sup> | Encaps.<br>MSCs | P-value<sup>##</sup> | P-value<sup>###</sup> |
| **Collagen**<br>**accumulation** | 0,43±0,049 | 0,097±0,01 | **<0,0001** | 0,077±0,01 | **<0,0001** | 0,94 |
| **S nucleus,**<br>**mkm<sup>2</sup>** | 21,7±5,6 | 44,2±9,7 | **<0,0001** | 43,4±5,4 | **<0,0001** | 0,93 |
| **S cell, mkm<sup>2</sup>** | 122±4,4 | 152,5±4,2 | **<0,0001** | 155,1±3,9 | **<0,0001** | 0,9 |
| **NC ratio** | 0,21±0,04 | 0,39±0,07 | **<0,0001** | 0,42±0,09 | **<0,0001** | 0,33 |

<sup>#</sup> p<0.05 – sham control vs. MSC administration
<sup>##</sup> p<0.05 – sham control vs. encapsulated MSC administration
<sup>###</sup> p<0.05 – MSC administration vs. encapsulated MSC administration

**Fig 6.**
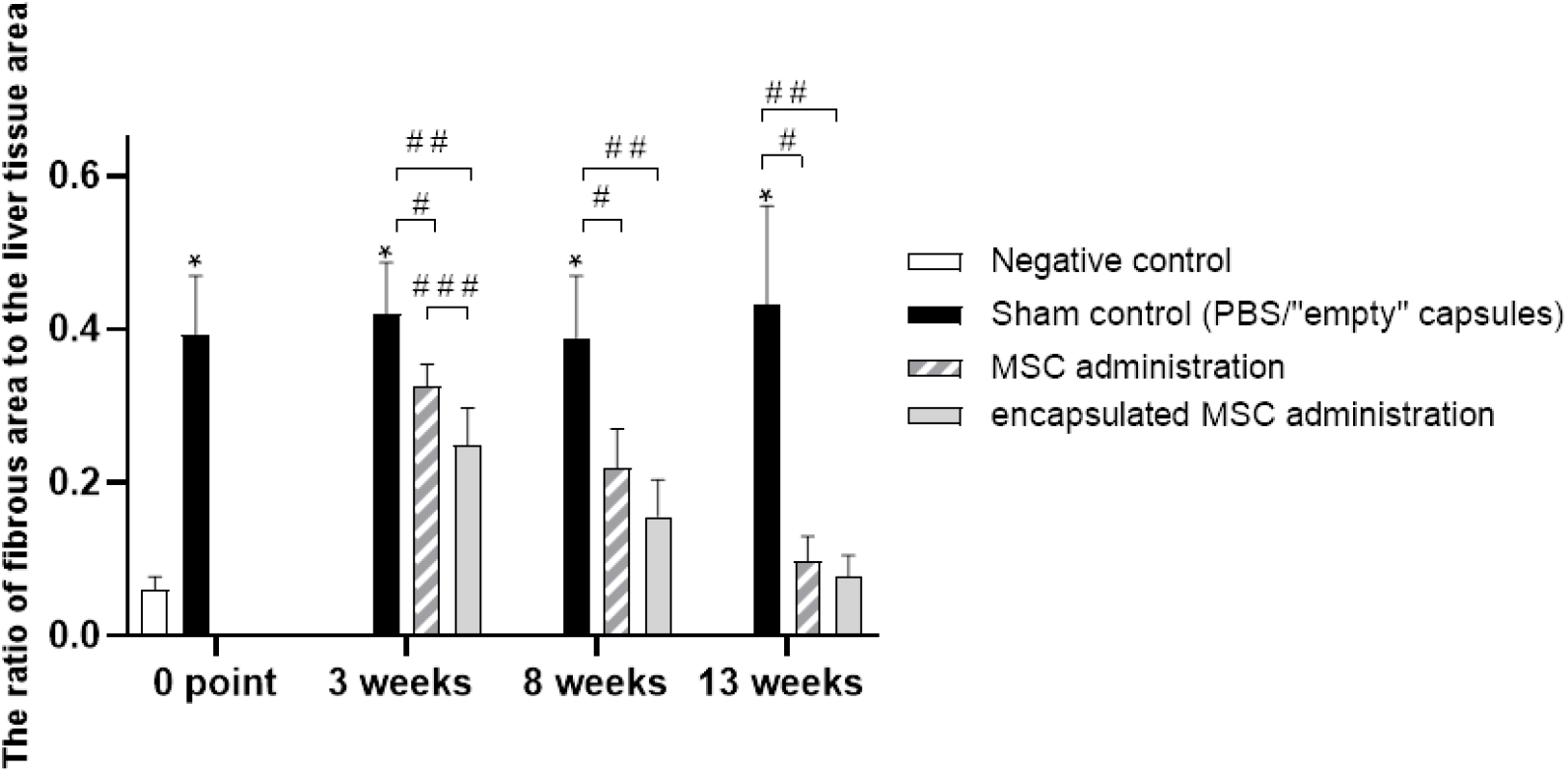
Effect of MSC transplantation on collagen accumulation in rat liver tissue: 0 point - 13 weeks after intraperitoneal administration of CCl_4_ to rats; 3, 8 and 13 weeks - after intravenous administration of MSCs and intraperitoneal administration of encapsulated MSCs to rats with liver cirrhosis. Results are expressed as mean±SEM. P values were calculated by ANOVA with Tukey’s test. There were no significant differences between the (sham control-PBS) and (sham control-«empty» capsules) – 0,1 < p < 0,9. ^*^ p<0.05 – negative control vs. sham control ^#^ p<0.05 – sham control vs. MSC administration ^##^ p<0.05 – sham control vs. encapsulated MSC administration ^###^ p<0.05 – MSC administration vs. encapsulated MSC administration

**Fig 7.**
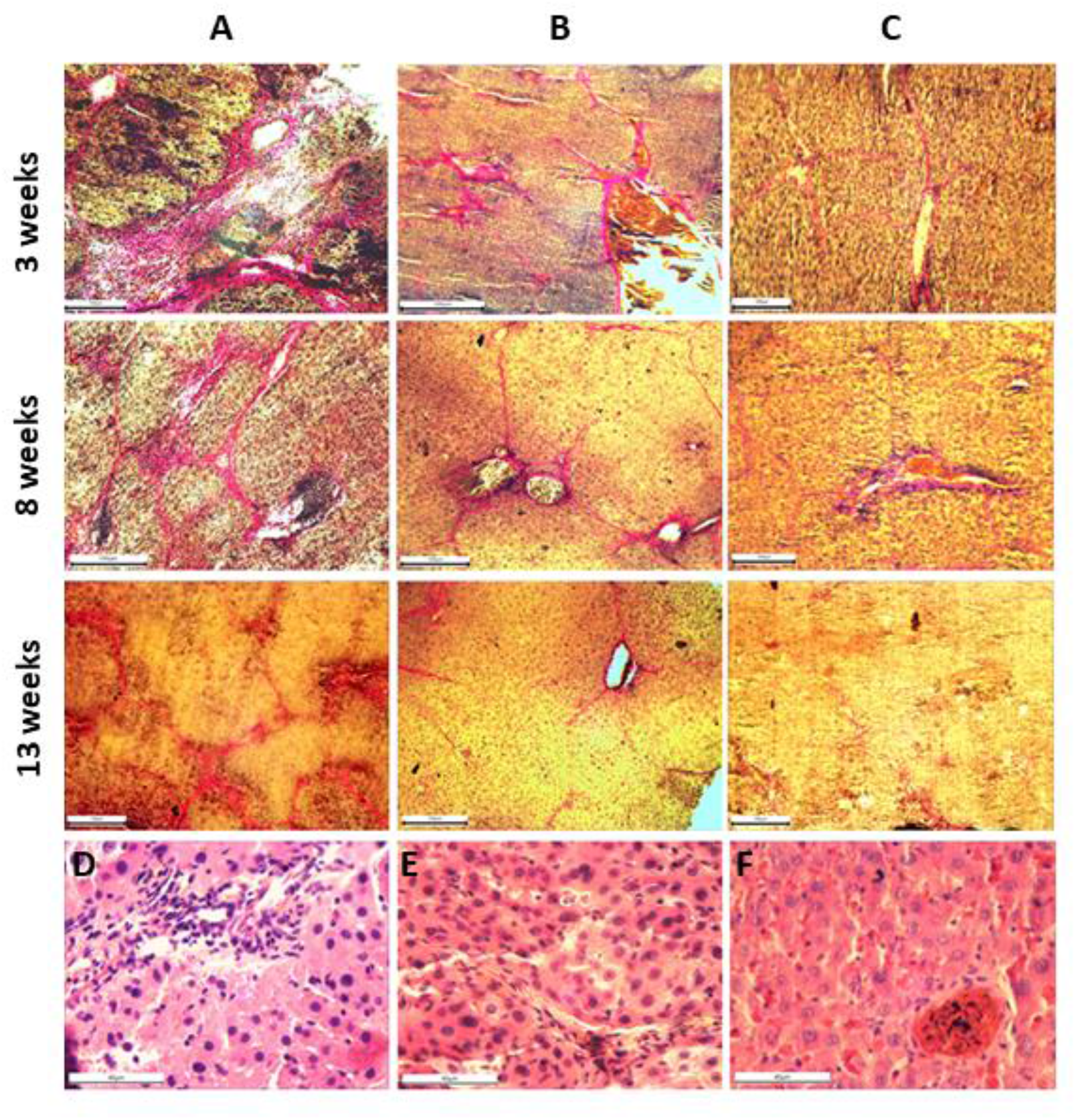
Histological analysis of liver tissue sections after hUC-MSC transplantation to rats with CCl_4_-induced liver fibrosis/cirrhosis. Van Giezon staining and H&E staining of rat liver sections 3, 8 and 13 weeks after MSCs transplantation: **A** - stained liver sections from a sham rats with liver cirrhosis which did not receive MSC transplantation; **B** - stained liver sections from rats which received intravenous injection of MSCs; **C** - stained liver sections from rats that received an intraperitoneal injection of encapsulated MSCs; Scale bar: A – C - 100μm, D – F - 40μm.

### The effect of MSC transplantation on the expression of certain genes associated with the liver fibrosis development

HUC-MSC transplantation led to a decrease in the level of EGF gene expression (Fig. 4), which had sharply increased during the formation of liver cirrhosis in rats. The decrease in EGF expression occurred faster in animals, which had received the encapsulated MSCs: three weeks after transplantation of encapsulated MSCs this gene was practically unexpressed, while after intravenous administration of the MSCs, EGF expression remained at a high level, almost disappearing up to 8^th^ week (Fig. 4A). Also by the 8^th^ week, α-SMA expression had returned to the level observed in the control animals, regardless of the transplantation route (Fig. 4B). GFAP expression decreased after 3 weeks almost to the level determined in animals that were not injected with CCl_4_ (Fig. 5A). The expression of eNOS, which decreased slightly during the development of cirrhosis (Fig. 5B), returned to control after 3 weeks in animals receiving any type of MSC treatment.

## Discussion

Mesenchymal stem cells exhibit some unique properties that make them a most promising object for use in cell therapy for various diseases, including liver diseases. In our study, we used human umbilical cord MSCs, because these cells have several advantages compared to MSCs from other tissues. They are an intermediate type between adult and embryonic stem cells. HUC-MSCs express pluripotency markers: NANOG, OCT-4, and SSEA-4, which are characteristic of embryonic stem cells [37]. Human umbilical cord MSCs have a higher level of expression of many genes compared, in particular, with bone marrow MSCs and this causes the unique therapeutic properties of these cells [38]. HUC-MSCs are hypoimmunogenic, since they practically do not express human leukocyte antigen D-related (HLA-DR) [39, 40]. For these reasons, they can be effectively applied for both allogeneic and xenogenic transplantation. In addition, hUC-MSC has a wider differentiation potential than MSC from other sources and differentiate, including hepatocyto-like cells [41, 42].

Our results on the transplantation of hUC-MSCs in rats with liver cirrhosis induced by CCl_4_ show that liver fibrosis disappears within 13 weeks after transplantation, and the main signs of the liver parenchyma normalize. The expression of genes associated with liver injury and the development of fibrosis also returns to the levels observed in negative control animals, both after systemic administration of hUC-MSCs and intraperitoneal administration of encapsulated MSCs. The recovery rate of parameters such as collagen accumulation and the level of EGF expression were higher with the transplantation of encapsulated MSCs. 3 weeks after transplantation of encapsulated MSCs, EGF expression was 3,4 times lower than after systemic administration of MSCs.

The use of different transplantation routes allows us to consider various mechanisms that determine the therapeutic effects of MSCs. Recent studies show that the mechanism that leads to the realization of the therapeutic potential of MSCs *in vivo* is not fully understood. The main factor providing the immunomodulatory properties of MSCs is considered their secretome. The use of MSC-conditioned culture media for therapeutic purposes [43-45], as well as enhancing of the MSC efficacy under the influence of MSC preconditioning by various factors that modify the spectrum of secreted molecules, confirm the decisive importance of the secretome for the therapeutic potential of MSCs [46-51]. In this study, two methods of MSC administration were used, which determine different mechanisms of influence on rat liver. Upon intravenous administration of MSCs, the main part of the cells is captured by the lungs and enough quickly disappear, however, a small portion of transplanted cells can achieve the liver [52-54]. Interestingly, the preconditioning of hUC-MSCs with some cytokines and other factors not only enhances immunomodulatory properties of MSCs but also affects their biodistribution, decreasing the amount of MSCs in the lungs and increasing in the liver [26, 55]. Thus, it remains unclear how the anti-inflammatory and regenerative effects of MSC in the damaged liver are implemented with such a fate of transplanted MSC, although the therapeutic effect is completely manifested. The immunomodulatory properties were found even in inactivated and apoptotic MSCs, which no longer secrete protein molecules, but at the same time activate immune system cells such as monocytes and macrophages due to MSC phagocytosis [56-59]. The liver repair, which is observed after the intraperitoneal administration of encapsulated MSCs, is caused only by the action of the factors secreted by them. Introduced into the blood MSCs quickly disappear from the body, probably as a result of phagocytosis by monocytes and macrophages. When encapsulated MSCs are administered intraperitoneally, the soluble factors secreted by them enter the abdominal cavity, where a large number of peritoneal macrophages are localized [60]. In our study, encapsulated MSCs secreted soluble factors for quite a long time, at least not less than one month (unpublished data).

Numerous studies show that under the influence of the factors secreted by MSCs macrophages polarize, transforming into an anti-inflammatory M2 phenotype, suppressing inflammation by increasing the expression of anti-inflammatory cytokine IL-10 and the scavenger receptors CD206 and CD163 [61-64]. It is also known that in response to liver injury, macrophages of the abdominal cavity rapidly invade into afflicted liver tissue [65]. Thus, the therapeutic effect of the intraperitoneal injection of encapsulated MSCs is obviously associated with the penetration of polarized macrophages into the liver, although polarization of hepatic macrophages is not excluded. The authors of the study, who injected MSCs intraperitoneally into mice with induced colitis, believe that the mitigation of colitis is associated with the polarization of tissue macrophages under the influence of chemokines, which can act on tissue macrophages at a distance. In our study, both routes of MSC transplantation led to almost complete liver recovery within 13 weeks, i.e. both delivery methods provided a high therapeutic effect of MSCs. At the same time, the rate of recovery of such parameters as collagen accumulation, some morphometric characteristics, and the level of EGF expression was higher with the transplantation of encapsulated MSCs. Apparently, earlier restoration of liver tissue is associated with the well - known advantages which acquire cells after encapsulation. These include protection from the effects of the immune system and increase the long-term survival of MSCs, which leads to the prolonged secretion of paracrine factors. [66-68]. Our results confirm that encapsulated MSCs have increased efficacy. Using different routes of MSCs administration on models of various diseases shows ambiguous results. Comparison of intravenous and intrasplenic administration of MSCs demonstrates the advantages of intravenous administration in the rat liver fibrosis model [69]. In another study, intraportal administration of MSCs is more effective than intravenous or intrahepatic and intraperitoneal [71]. The study of the efficiency of four methods of MSC transplantation for the treatment of acute liver damage in rats showed no significant differences between injections in the hepatic artery, the portal vein, and the caudal vena [72]. The therapeutic effect was not observed with an intraperitoneal injection of BM-MSCs which is contrary to our results with the intraperitoneal delivery of encapsulated hUC-MSCs.

In a model of acute colitis in mice, it was shown that intraperitoneal and subcutaneous routes of MSCs delivery were more effective than intravenous route [73, 74]. Often, systemic and local administration of MSCs demonstrated nearly the same therapeutic efficacy, although when administered locally, MSCs survive longer. [75]. The study of the survival time of MSCs with different routes of transplantation showed that MSCs administered systemically were not detected after a few days, intraperitoneal and subcutaneous administration ensured the survival of MSCs for 3-4 weeks, and after intramuscular administration, the cells survived for more than 5 months [76]. This makes the intramuscular administration of MSCs a promising alternative to intravenous administration, in particular, for systemic diseases [77]. Thus, the results of numerous studies show that the MSC efficacy depends on the route of MSC delivery, but, apparently, it will not be universal for all diseases. In conclusion, in this study, we compared the therapeutic effect of systemic transplantation of human umbilical cord MSCs and intraperitoneal administration of MSCs encapsulated in alginate capsules in a CCl_4_-induced model of cirrhosis in rats. Transplantation of hUC-MSCs into rats with liver cirrhosis led to liver recovery both with systemic and with intraperitoneal administration of encapsulated MSCs. By the 13^th^ week after transplantation, fibrosis gradually disappeared, the expression of EGF, alfa-SMA, GFAP, eNOS was restored to the level characteristic of control animals that did not receive CCl_4_. The rate of improvement of the injured liver tissue is higher after transplantation of encapsulated hUC-MSCs. Thus, human umbilical cord MSCs demonstrate high therapeutic potential in liver recovery, both routes of MSC transplantation led to almost complete liver recovery within 13 weeks, but transplantation of encapsulated cells leads to a higher rate of liver recovery.

## Acknowledgments

We thank the Dr Geert Potters (Antwerp Maritime Academy, Belgium) made invaluable comments on language and style.

